# Quantal PCR: A Calibrator-Free Method for Determining the Unit for Nucleic Acid Quantification

**DOI:** 10.64898/2026.02.08.704629

**Authors:** Xiangrong Tang, Yunqing Wen, Rong Qin, Jishen Zhang, Zhijun Tang, Chenxiao Ding, Yan Zhang, Yang Tong

**Affiliations:** Jiangsu Precision Biotechnology Co., Ltd., Room 803, Building 9, Hengsheng Zhigu Technology Park, Xuzhou, Jiangsu, 221132, P. R. China; Center for Excellence in Brain Science and Intelligence Technology (Institute of Neuroscience), Chinese Academy of Sciences, 320 Yue Yang Road, Shanghai, 200031 P. R. China; Shanghai Hongshi Medical Technology Co., Ltd. 5th Floor, Building 5, No. 1181 Zhaoxian Road, Shanghai, 200182, P. R. China; School of Life Sciences and Biotechnology, Shanghai Jiao Tong University, 800 Dongchuan Road, Shanghai City, 200240, P. R. China; School of Pharmaceutical Science and Technology, Tianjin University, 92 Weijin Road, Tianjin, 300072, P. R. China

## Abstract

Quantitative polymerase chain reaction (qPCR) is limited in measuring absolute nucleic acid copy numbers due to the inherent variability of calibrators. Here, we introduce the Quantal PCR (quPCR), a novel method that eliminates the need for calibrators by defining an intrinsic quantal unit derived from the thermodynamic and kinetic properties of the replication system. This approach first determines amplification efficiency at high template concentrations, which is then used as the replication probability to construct quantification cycle (Cq) distribution profiles. These profiles are compared with those from limiting dilution PCR to derive the Cq value for the minimal quantal-replication unit (“quCq”), enabling calculation of the sample copy number. Validation using a dual-target DNA template showed near-identical copy numbers using two distinct target-specific replication systems. Thus, quPCR represents a new method for absolute nucleic acid quantification at the single-molecule level, offering a calibrator-free alternative for absolute quantification.

---

Real-time quantitative PCR quantifies nucleic acids by comparing the fluorescence-based quantification cycle (Cq) values of samples to calibrated standards^1^, yet its accuracy is limited by the calibrators’ concentration measurements (e.g., UV absorbance), which are susceptible to interferences^2^. Digital PCR provides absolute quantification without calibration by partitioning samples and applying Poisson statistics^3^, but it faces constraints including a limited dynamic range, partition volume variability^4^, and potential deviation from Poisson distribution due to template aggregation^5^.

Relative quantification presents another qPCR strategy, based on the following equation^6^, where the ratio of sample copy numbers D is given:

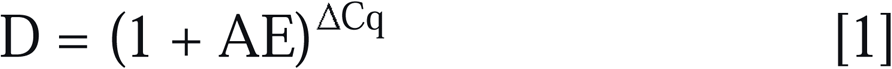

where ΔCq represents the difference of Cq at two different concentrations, and amplification efficiency (AE) is defined as the fraction of template molecules copied in each cycle^7^. However, it lacks an absolute reference unit and AE is usually not exactly 100%, thus failing to provide accurate absolute quantification. Building upon the relative quantification framework, we here developed an approach that precisely determines AE and replaces the arbitrary calibrator with a quantal unit—the smallest indivisible entity that represents a single-copy ssDNA molecule. We define the quantal-replication unit Cq (quCq) as the theoretical replication cycle number at which replication based on a single molecule reaches the fluorescence threshold. Substituting quCq into the relative quantification equation 1 yields the equation for the absolute quantification of sample copy number Ns:

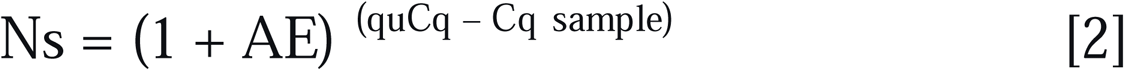

At the molecular level, amplification depends on the probability of a single molecule being replicated—a stochastic process. In this study, we determined the AE —conceptualized as the per-cycle replication probability—from the ΔCq between two high-concentration samples. This probability, as an intrinsic property of the replication system, exhibited remarkably low variability, providing a stable foundation for extrapolation to the single-molecule level. The quCq was then determined by matching the dominant peak (mode) of the experimental Cq distribution from a limiting dilution assay to the simulated profile for a single molecule.

The reliability of this method was tested using a dual-target DNA template, where near-identical copy numbers were obtained from two independent replication systems. Here, we present Quantal PCR (quPCR), a method establishes a framework for calibrator-free absolute quantification at the single-molecule level, with broad potential utility in research and diagnostics.

We first sought to precisely determine the amplification efficiency (AE). In a dilution experiment, the dilution factor (D) represents the ratio of concentrations before and after dilution, and ΔCq denotes the difference in Cq values between two different dilutions. We first transformed the equation [1] into the following form:

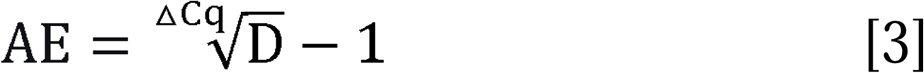

To illustrate the determination of the amplification efficiency AE, we performed qPCR for the RNase P gene fragment (hereafter termed the “R system”, Table S1). A 50-fold dilution of the sample containing the fragment was used to determine Cq at two different concentrations. We performed 10 independent experiments for this pair of template concentrations, each with 48 qPCR replicates (Figure 1A). The AE was calculated as 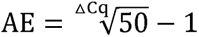, yielding a mean value of 0.959 ± 0.006 (SEM, n = 10). Nearly identical mean AE values were found when a 100-fold dilution (D = 100) or a series of 5- and 10-fold dilutions were used (Figure 1B). Our method yielded significantly lower standard deviation than the traditional calibration curve approach. In this method, the AE detected across different dilution factors remains stable; we therefore postulate that AE is a stable intrinsic property of the replication system under stable thermodynamic and kinetic conditions. Consequently, to determine AE accurately, it is essential to minimize the impact of two major stochastic error sources prevalent at low copy numbers (<10^4^): binomial sampling error during dilution (Table S2) and the probabilistic “all-or-none” initiation of replication in early cycles (Figure S1, as described below).

**Figure 1.**
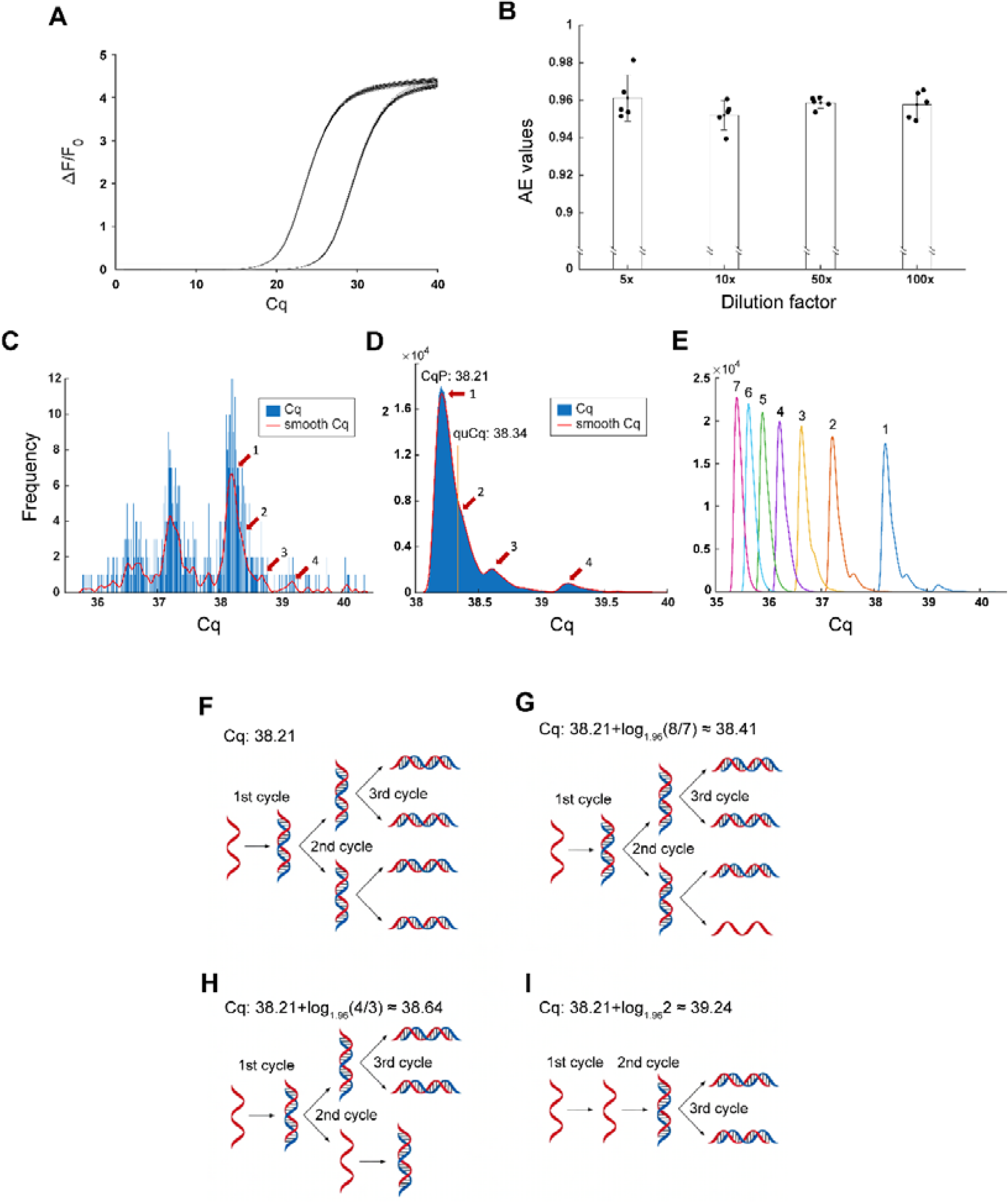
Amplification efficiency (AE) determination. **A)** Fluorescence curves from a 96-well plate containing ∼10^8^ copies/mL template and its 50-fold dilution (48 wells each). **B)** The AEs were measured for each dilution factor with 5 experimental replicates, which showed no statistically significant differences among the groups (pooled *t*-test). **C)** The Cq distribution from 960 qPCR assays of limiting dilution of ssDNA, with Gaussian smoothing shown in red. Peaks 1 – 4 correspond to those in panel D. **D)** Simulation of the Cq distribution for a single ssDNA template copy. The red curve shows the Gaussian-smoothed result. **E)** Cq distributions for 1 – 7 initial templates (numbered 1 – 7). **F-I)** Mechanism underlying the main peaks in the Cq distribution profile for a single-copy template. Panels F, G, H, and I show the formation of peak 1, peak 2, peak 3, and peak 4 in panels C and D, respectively.

Next, we employed a limiting dilution assay to determine the Cq value for replication at extremely low template concentrations of ssDNA of the RNase P gene fragment, in order to identify the potential quantal unit of replication. We next performed replication of ssDNA samples at limiting dilution in 96-well plates at the concentration of ∼1/3 of the wells were tested negative, based on the Poisson distribution at the extremely low concentration (Figure S2, Table S3). A total of 960 qPCR assays were performed across ten 96-well plates, and Cq values of positive reactions were plotted, and Gaussian smoothing was used to help the identification of the peaks in the distribution patterns (Figure 1C).

Recognizing the quantized nature of nucleic acid molecules, we developed a discrete mathematical model to simulate the amplification kinetics of the R system at various concentrations. Under the condition of AE = 0.959, we simulated the probabilistic replication process of each template molecule for specific starting copy numbers until amplification reached (1+AE)^20^ copies. Starting with a single template, 10^6^ replication simulations were performed (Figure 1D). The threshold for all simulations was set to a copy number of (1+AE)^quCq^, where the quantal-replication unit Cq (quCq = 38.34) represents the theoretical Cq value for a single copy, as derived from the peak-matching calibration described below. The threshold replication copy (1+0.959)^38.34^ was then applied to the simulations for different initial template numbers. The results of our simulations revealed skewed, multimodal Cq distributions, with a narrow and sharp dominant peak, and with increasing skewness and multimodality at lower template concentrations (Figure S1). The apparent multimodal distribution in the Cq profile originates from the quantized replication of the template. Figure 1F-I illustrates the probabilistic replication process for the single initial template that yielded the four predominant peaks in the distribution profile in Figure 1C, D: The highest peak corresponded to reactions in which all template molecules are replicated in all first three cycles (Figure 1F), whereas the 2^nd^, 3^rd^, and 4^th^ peak reflected a fraction (1-AE, or ∼4%) replication failure in the 3^rd^, 2^nd^, and 1^st^ cycle of replication (Figure 1G-I). The Cq values of the three highest peaks obtained by simulations aligned well the profiles of the experimental results. This analysis showed that quantal analysis at the limiting dilution allows the determination of quantal unit of Cq.

Using the peak value of Cq distribution (CqP = 38.21) and the measured AE (0.959 ± 0.006) as inputs, we determined the quantal-replication unit Cq (quCq) via stochastic modeling. The model iteratively refined a candidate quCq value until the simulated dominant CqP aligned with the experimental CqP, yielding a final quCq of 38.34. This fundamental parameter represents the theoretical Cq expected for a single molecule to reach the detection threshold under the system’s fixed AE, enabling calibrator-free quantification via:

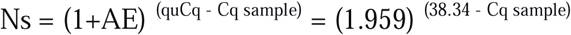

We also simulated Cq distribution profiles for initial copy numbers from 1 to 7 (Figure 1E) and determined boundary values. This allowed us to “count” the initial copy number in each well based on its Cq value (Table S4). The sum of copy numbers per plate was consistent with Poisson estimates from negative wells (Figure S3). We also observed a few reactions with Cq values higher than that of peak 4 (Figure S4). We hypothesized that 50 cycles of PCR followed by 80 cycles of linear PCR on a ∼300-base template introduced stochastic mutations in primer or probe binding sites, causing delayed Cq. Sequencing of these deviated products confirmed mutations in the probe binding site (Figure S5).

The entire methodology was also applied to the qPCR system for the HER2 gene fragment (hereafter referred to as the “H system”, Table S1). The skewed multimodal Cq distribution and stochastic amplification nature were reproduced, confirming the general applicability of quPCR. An AE of 0.954 ± 0.006 and a quCq of 38.64 were determined (Figure S6).

To assess the accuracy of the quPCR method, we performed a ratio recovery experiment on a synthetic DNA template with a predefined 1:1 stoichiometry of two replication targets (RNase P and HER2) across five concentrations, using two different replication systems. The measured ratios were consistently close to 1 (1.022 ± 0.035, SEM, n = 5; Table S5), demonstrating the high accuracy and robustness of the quPCR method in the absence of an absolute certified standard.

A fundamental problem of conventional qPCR is that its accuracy hinges upon external calibrators. We surmised that AE is an intrinsic property of the replication system. It remains constant over template concentration corresponding to Cq of 18 to 25, within the linear dynamic range of replication where the depletion of substrate (e.g., primers, dNTPs) and temperature-dependent changes in template structure distributions is negligible. Thus, a relatively high initial template concentration could be used to calculate AE, which can then be used to estimate the quantal unit of replication via the determination of quCq.

Once the low- and high-concentration samples reach equal template copy numbers during replication, their subsequent replication kinetics converge. Therefore, a change in amplification efficiency (AE) by the time the Cq threshold is reached does not affect ΔCq measurement. Circular or long nucleic acid templates may lower early-cycle amplification efficiency due to secondary structures, potentially underestimating initial copy numbers^8,9^. However, this structural effect diminishes in later cycles, where efficiency is determined by the shorter amplicon. Moreover, when such templates are used in a calibration curve, their structure systematically affects Cq values across all dilutions, and this effect is canceled out in ΔCq-based AE calculations.

With determining the AE, we anchored the Cq scale to the single molecule-the fundamental quantal unit - through limiting dilution and stochastic modeling. This was achieved via a “peak-matching” strategy, where the sharp dominant peak of the experimental Cq distribution was matched to the simulated profile for a single molecule. This confers the dual practical benefits of reducing the burden of experimental replication and providing robustness against outlier data points. The resulting quantification equation, Ns = (1 + AE)^(quCq - Cq sample)^, represents a powerful differential measurement. Here, the quCq constant encapsulates and cancels out the complex kinetics of amplifying a single target molecule to the fluorescence threshold, through its comparison with the sample Cq.

Furthermore, quPCR redefines key metrological concepts of qPCR. It establishes the single molecule as the unambiguous limit of detection (LOD)^10^. Positivity is determined not by an arbitrary Cq cutoff but by the characteristic Cq distribution of this minimal unit, inherently accounting for the stochasticity of amplification.

Our quPCR approach represents a fundamental reconceptualization of nucleic acid quantification. It achieves a paradigm shift by replacing dependence on external calibrators with an intrinsic, self-calibrating framework based on system-specific constants, thus redefining the core principles of measurement itself without introducing new reagents or instruments. We anticipate this accessible yet powerful framework will be applicable for many precise molecular measurements from nucleic acids to proteins and beyond.

## ACKNOWLEDGMENTS

We are profoundly grateful to Dr. Mu-ming Poo for his insightful critique of the manuscript, his decisive editorial guidance in shaping the final version, and his invaluable support in championing this work. His mentorship was pivotal to this research. We thank for Drs. Shiqing Cai, Hao Wu, Xiaodong Yang, Yu Zhou, Mengwei Huang for suggestions.

## Author Contributions

X.T. and Y.W. designed the study; X.T., J.Z., Z.T. and C.D. performed the experiments; Y.W. and R.Q. wrote the data processing program; X.T., Y.W., Q.R., J.Z., Z.T., C.D., Y.Z., and Y.T. participated in the data curation; and X.T. and Y.W. wrote the manuscript.

## Supplemental Information

### Materials and Methods

#### Oligos, Reagents and Equipment

The templates used in this study were synthesized by Map Biotech (Shanghai, China). Primers and probes were synthesized by Takara Bio (Dalian, China). All sequences are provided in Table S3. Other qPCR reagents including the reaction buffer, dNTP mixture, TaqHS DNA polymerase were purchased from Takara and the Low-adhesion qPCR 96-well plates and pipette tips were purchased from Corning (NY, USA). All qPCR experiments were performed on a SLAN-96S real-time qPCR machine from Hongshi Medical (Shanghai), with the relative fluorescence threshold set at the instrument-recommended value of 0.12 unless otherwise specified.

#### qPCR and linear PCR protocols

All PCR amplifications were performed in a 25 μL total volume containing: 2.5 μL PCR Buffer (Takara), 1 μL of 2.5 mM dNTP mixture, 1 μL of TaqHS (1 U/μL), and 5 μL of template. The standard thermal profile was: initial denaturation at 95 °C for 10 min, followed by cycles of 30 s at 95 °C and 60 s at 60 °C, unless otherwise specified. The specific composition and any modifications for each application are detailed below:

qPCR reactions for the RNase P (R) gene system contained 0.2 μL of the TaqMan probe, 0.4 μL each of the 10 μM forward and reverse primers, and the annealing/elongation step was extended to 90 seconds. For the HER2 (H) gene system, reactions contained 0.2 μL of the TaqMan probe and 1 μL each of the 10 μM forward and reverse primers.

Template Preparation (for both R and H systems):

Regular PCR (dsDNA template): Reactions contained 0.4 μL each of the relevant 10 μM forward and reverse primers.

Linear PCR (ssDNA template): Reactions contained 0.04 μL of the relevant 10 μM reverse primer only (no forward primer).

#### AE determination

For the measurement of Amplification Efficiency (AE), synthetic DNA fragments encompassing the target regions of the human RNase P gene (for the *R* system) or the HER2 gene (for the *H* system) were synthesized and cloned into the pUC57 plasmid. These constructs served as the defined templates for subsequent AE determination experiments.

The AE for each qPCR system was determined within the linear dynamic range of replication (Cq 18–25), which corresponds to initial template concentrations of approximately 10^6^ to 10^4^ copies per reaction as established by quPCR.

AE was assessed using multiple, independent dilution schemes for both systems. For the *R* system, the following were prepared from a high-concentration stock (∼10^8^ copies/mL, Cq ≈ 18): (i) single-step 50-fold and 100-fold dilutions, and (ii) multi-step 5-fold (three steps) and 10-fold (two steps) serial dilutions. For the *H* system, the single-step 50-fold dilution was performed. To control sampling error at this initial stage, each dilution scheme was performed with a consistent number of 5 independent replicates, generating independent template of different dilutions for qPCR. Although the mean amplification efficiency (AE) values measured at different dilution factors showed no significant differences (Figure 1B), the 50-fold dilution offers greater operational convenience.

In the qPCR assay, each template concentration from the above dilutions was first thoroughly mixed with a master mix. This homogeneous pre-mixture was then aliquoted into a defined number of technical qPCR replicates.

The minimum number of replicates (n) required to ensure with > 99% probability that the sample mean deviates by less than a defined tolerance (*δ*) from the expected value (μ *= N·p*) is derived from the following condition (based on binomial distribution):

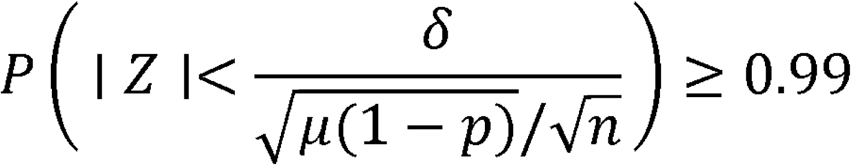

Where *Z*_O.995_ = 2.576, *N* is the total template number in the master mix aliquot, and *p* is the sampling probability.

For the 50-fold dilution of the *R* ystem, using a stock with *N* = 500,000 molecules and setting the tolerance *δ* = 100 copies from the expected *μ* = 10,000, solving the equation yields n ≥ 6.5. Therefore, at least 7 technical replicates are required to meet this precision goal. Our experimental design used 48 replicates for this concentration point, substantially exceeding the statistical minimum to ensure sampling error was rendered negligible.

For all other dilution schemes (100-fold, 5-fold series, 10-fold series for *R*, and 50-fold for *H*), the required minimum number of technical replicates (n) for each concentration point was similarly calculated based on its specific *N*, p, and precision goal (*δ*). The actual number of replicates used (e.g., 48 for single-step dilutions, 16 or 24 for steps within serial dilutions) was chosen to meet or far exceed the respective calculated minimum in each case, thereby robustly controlling sampling variability across the entire study.

The method for calculating AE depended on the dilution scheme:

- For single-step dilutions, the ΔCq for each independent dilution replicate was calculated and applied to the formula: AE **=** D^1/ΔCq^ - 1, where D is the dilution factor.
- For multi-step serial dilutions, AE was derived from the slope of the linear regression of Cq values plotted against the logarithm (base D) of the dilution factor across all steps within that series.

The final reported AE for each system and dilution scheme represents the mean and standard error calculated across all independent dilution replicates. This statistically-grounded approach ensures that the derived AE is a precise and stable estimator of the system’s intrinsic replication property.

#### Cq distribution profiles at limiting dilution and simulated quantal analysis

In the limiting dilution assay, a total of ten 96-well plates were used, and the Cq values from all positive replicates were employed to construct the Cq distribution profile. The distribution was divided into 1800 equally spaced bins, as shown in Figure 1C. Gaussian smoothing with a sliding window width of 0.15 cycles was then applied to get the low noise distribution (red line in Figure 1C). The most prominent peak (marked with 1) corresponding to all ssDNA amplified in the first three cycles (Figure 1F) and three other major peaks (marked with 2 - 4) corresponding to one ssDNA not amplified in the first 3 cycles (Figure1 G - I) were observable.

A model for extraction of the Cq distribution profiles was established. Briefly, in cycle 1 - 20, the stochastic nature of qPCR was fully considered when judging the AE-dependent amplification of each ssDNA molecule. A random value in the range of 0-1 was generated for each copy of ssDNA at the beginning of these cycles. When this value ≤ AE (0.959), an amplification event was called. When this value > AE, failure in amplification was called. The copy number of templates in each reaction can be calculated cycle by cycle. The copy number at the end of cycle 20 was determined to be N_20_. This process was repeated 1× 10^6^ times to generate 1× 10^6^ N_20_ values. To accelerate the modeling, starting from the 21^st^ cycle, where contribution to the Cq variation from individual ssDNA templates is much less significant, the threshold copy number N_quCq_ = (1+AE) ^quCq^ = 1.959^38.34^ was used to determine 1 × 10^6^ possible Cq values with the formula, Cq = lg(N_quCq_ /N_20_) / lg(AE+1) + 20 – 1 = lg(1.959^38.34^ /N_20_) / lg(1.959) + 19. It was noted that 38.34, a value obtained in a practical system (see below) was given as the quCq value. The distribution histogram was plotted with the Cq data divided into 1000 equal-width bins, as shown in Figure 1D. Gaussian smoothing with a width equal to one-tenth of the 90% data range was applied to remove noise (red line in Figure 1D). The peaks labeled with numbers 1 – 4 in Figure 1D correspond to the four scenarios illustrated in Figures 1F–I. Similar processes were carried out for 1 – 7 ssDNA templates in Figure 1E and 2 – 10000 ssDNA templates in Figure S1. A symmetric, approximately Gaussian distribution emerged at initial template numbers above ∼100 molecules (Figure S1F-H). The spread of the Cq distribution profile also narrowed progressively with increasing template number. The scripts for this modeling are provided in the supplementary data.

#### Linear PCR to generate ssDNA template

An insert containing an RNase P, Gastrin and HER2 gene fragment in tandem were synthesized and cloned into the pUC57 vector, forming the plasmid pUC57RGH. This plasmid was used as template with primer pairs 1F-1 and 1R-1 for PCR amplification of linear dsDNA containing the RNase P fragment, and with primer pairs 2F-1 and 2R-1 for dsDNA containing the HER2 fragment, as shown in Table S3. These two PCR products were then diluted and used as template for linear PCR to generate ssDNA using a unidirectional primer 1F-1 and 2F-1 respectively. The linear PCR product was then subjected to limiting dilution for qPCR assays.

#### Comparison of quPCR quantitation with Poisson distribution

We modeled the Cq distribution profiles for initial copy number from 1 to 7 as described above (Figure 1E), and determined these boundary values. The boundaries of copy number are defined by the intersection points of the distribution outlines. These values, along with the false alarm and false rejection rates, are listed in Table S4 and used for the determination of the initial copy numbers in each of the qPCR assays following limiting dilution, which were also compared with Poisson distribution estimation (Figure S3, Table S2).

#### Ratio determination between binary targets

DNA fragments containing the RNase P and HER2 genes were synthesized and cloned into a pUC57 vector. The resulting plasmid was used as a template for PCR amplification of a 327 bp fragment containing both targets (Table S3). This PCR product was then diluted to roughly 10^8^ copies/mL, approximately corresponding to a Cq of 19, A serial five-fold dilution with TE buffer (1 mL sample into 4 mL TE) was performed to generate quPCR templates. quPCR quantitation of both targets upon measuring their AE and quCq values were carried out for each concentration. The ratio of the two targets at each dilution was compared to a theoretical 1:1 value and standard deviation was calculated (Table S5).

---

**Figure S1.**
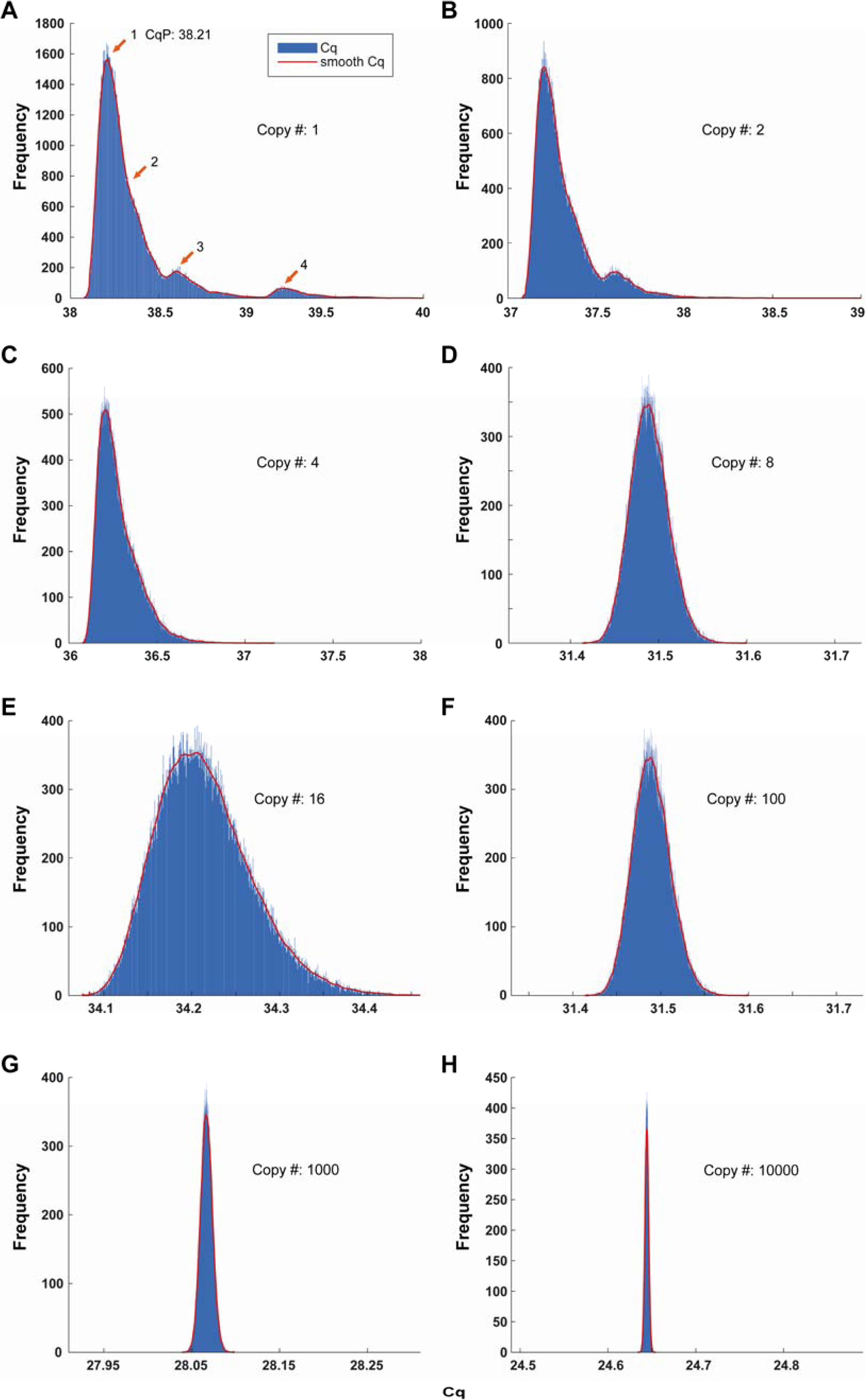
Computational simulation for the Cq distribution profiles of 100,000 qPCR assays with a ssDNA template. **A-D)** Initial copy numbers of the ssDNA template: 1, 2, 4, and 8 respectively; **E-H)** Initial copy numbers of the ssDNA template: 16, 100, 1000, and 10000 respectively. With increasing template copy number, the Cq distribution narrowed progressively, such that beyond 10,000 copies, the stochastic fluctuation arising from quantal replication was restricted to a range of less than 0.02 (Cq value).

**Figure S2.**
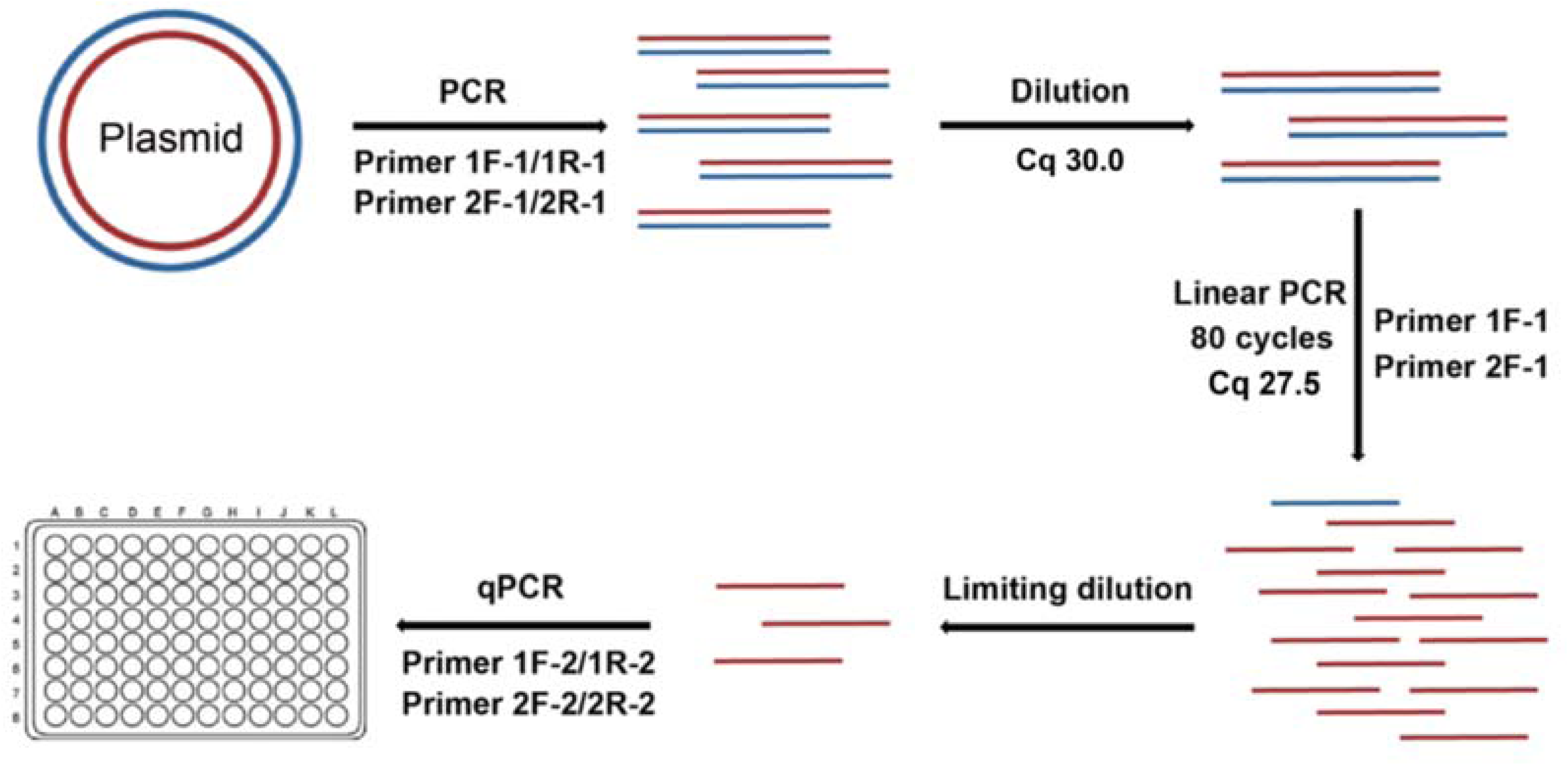
Schematic diagram showing template preparation for qPCR assays with a single copy ssDNA by linear PCR and limiting dilution. The plasmid pUC57RGH was used as template with primer pairs 1F-1 and 1R-1 for PCR amplification of linear dsDNA fragment containing an RNase P fragment, while the plasmid pUC57RGH was used as template with primer pairs 2F-1 and 2R-1 for dsDNA fragment containing a fragment. These two PCR products were then diluted and used as template for linear PCR to generate ssDNA using a unidirectional primer 1F-1 and 2F-1 respectively. The linear PCR product was then subjected to limiting dilution for quPCR assays.

**Figure S3.**
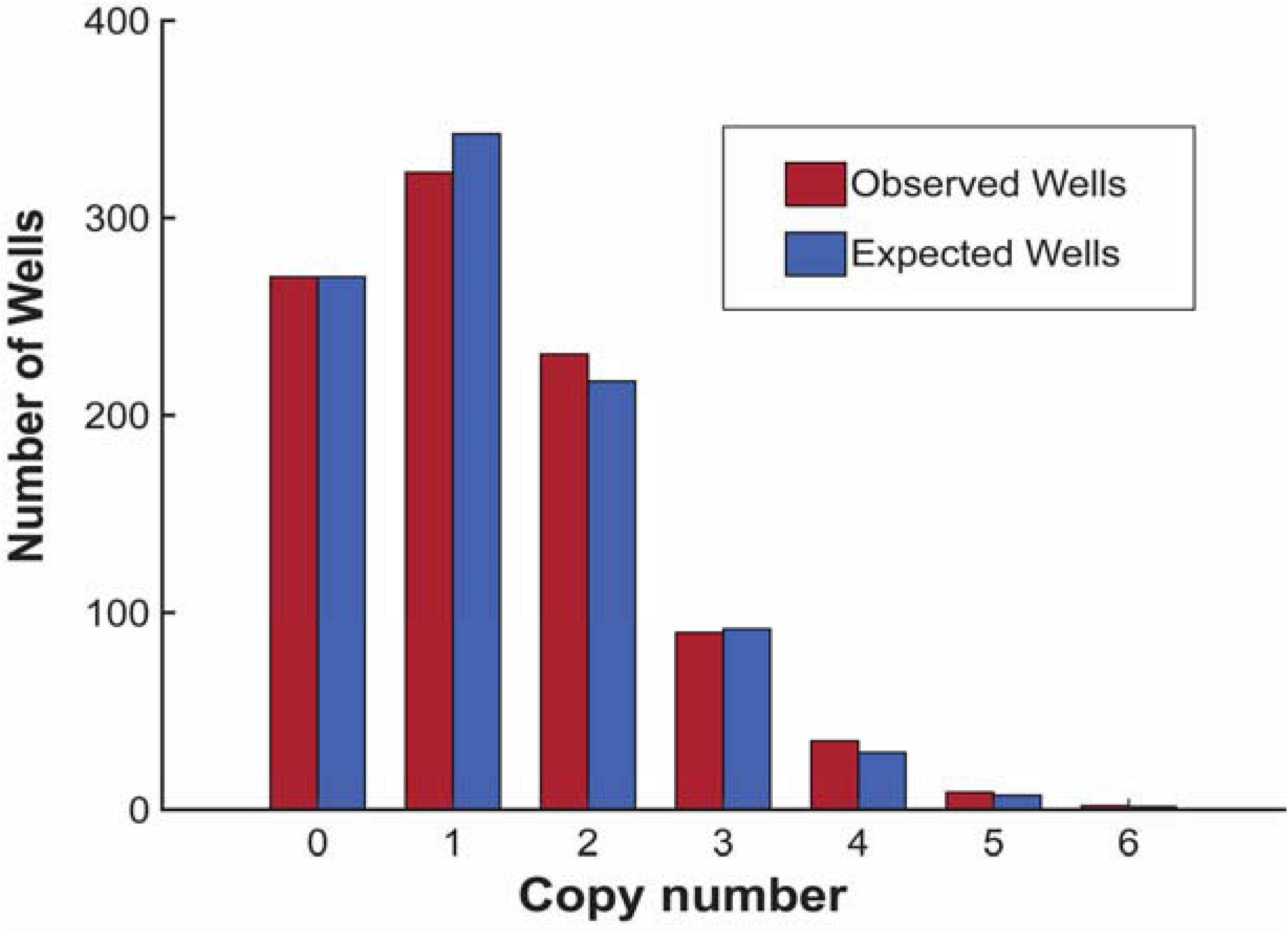
Comparison between quPCR and Poisson distribution estimation. The observed replicate well data (red) were obtained from ten 96-well plates, with the number of templates per well ranging from 0 to 6. Of the 960 replicate wells, 270 were negative, while the remaining wells contained a total of 1,252 templates. The average copy number per well was calculated based on the proportion of negative wells, and equal to -ln(270/960) = 1.269. The expected numbers of wells containing 0 to 6 templates were then derived from the Poisson distribution (blue). The chi-square goodness-of-fit test results (χ²(5) = 3.677, p = 0.597) indicate no significant deviation of the experimental data distribution from the Poisson distribution. The null hypothesis that the data follow a Poisson distribution cannot be rejected at the α = 0.05 significance level.

**Figure S4.**
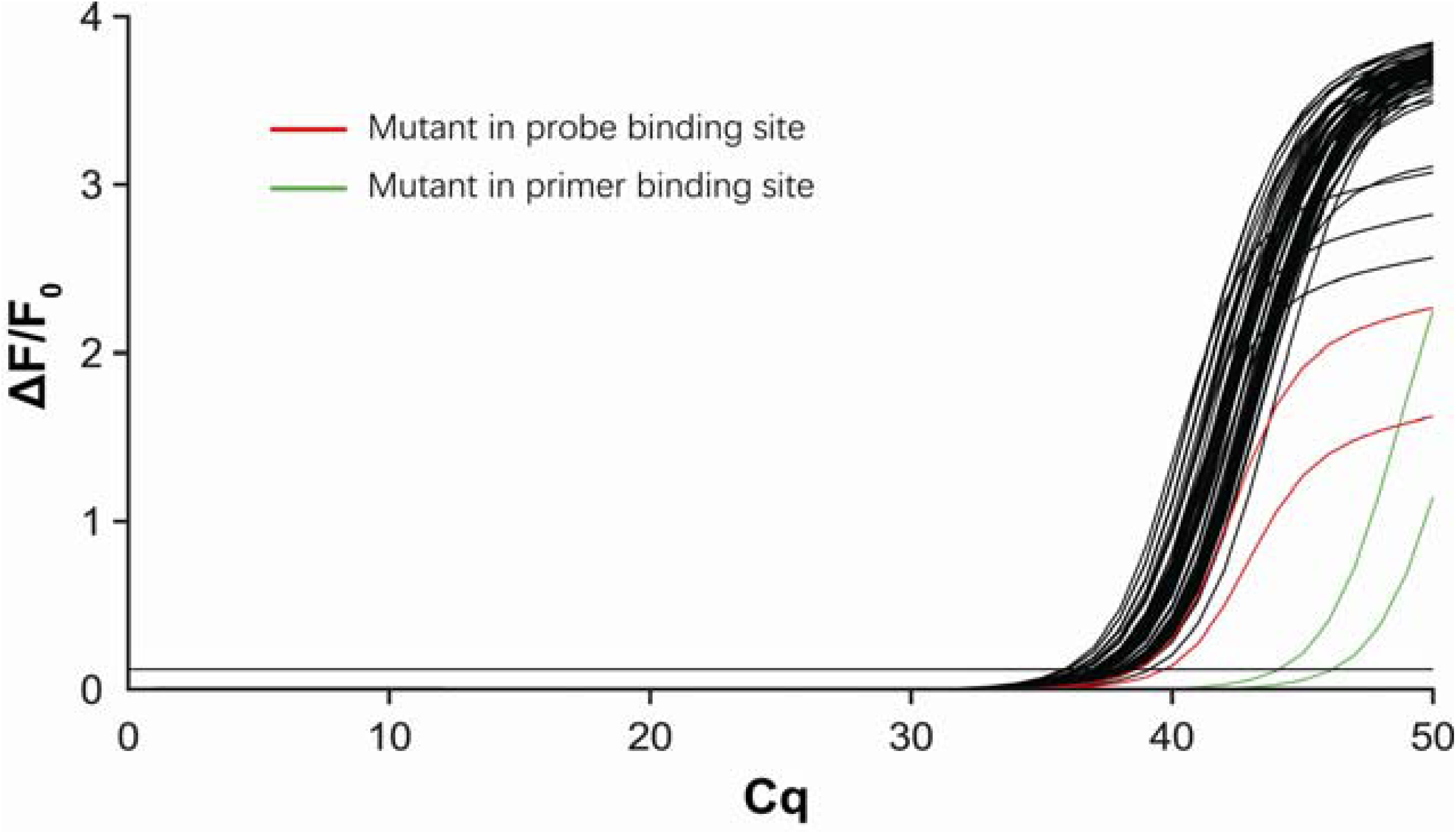
Fluorescence curves of qPCR assays in a representative 96-plate with ssDNA template limiting dilution analysis. Interestingly, we also observed a few qPCR fluorescence curves with Cq values higher than peak #4 (Figure 1C, D). This frequency might higher than the expected frequency of amplification abortion in multiple early cycles based on the known AE value. The fidelity of Taq polymerase used in our assays was reported to be as high as 0.01%, and could be compromised with increasing PCR cycles^1, 2^. Fifty cycles of PCR followed by 80 cycles of linear PCR with a template of ∼100 bases long are expected to introduce stochastic mutation in each 96-well plate, which, when occurs at either the primer or probe binding site, will result in a deviated Cq value. Cloning the qPCR product followed by sequencing revealed mutations in the probe binding site (Figure S5). By contrast, a mutation occurs at the primer binding site is not detectable by sequencing the PCR product.

**Figure S5.**
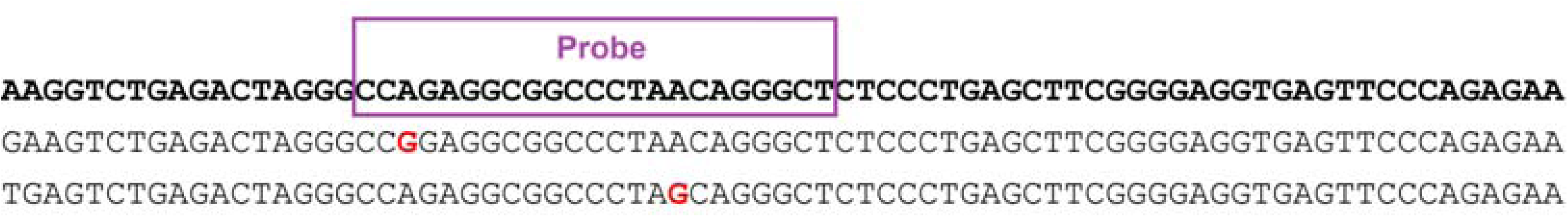
Sequencing results of the qPCR products in assays with deviated Cq values revealing errors in the probe binding area. The mutated base(s) within the probe region of the two PCR products are highlighted with red characters.

**Figure S6.**
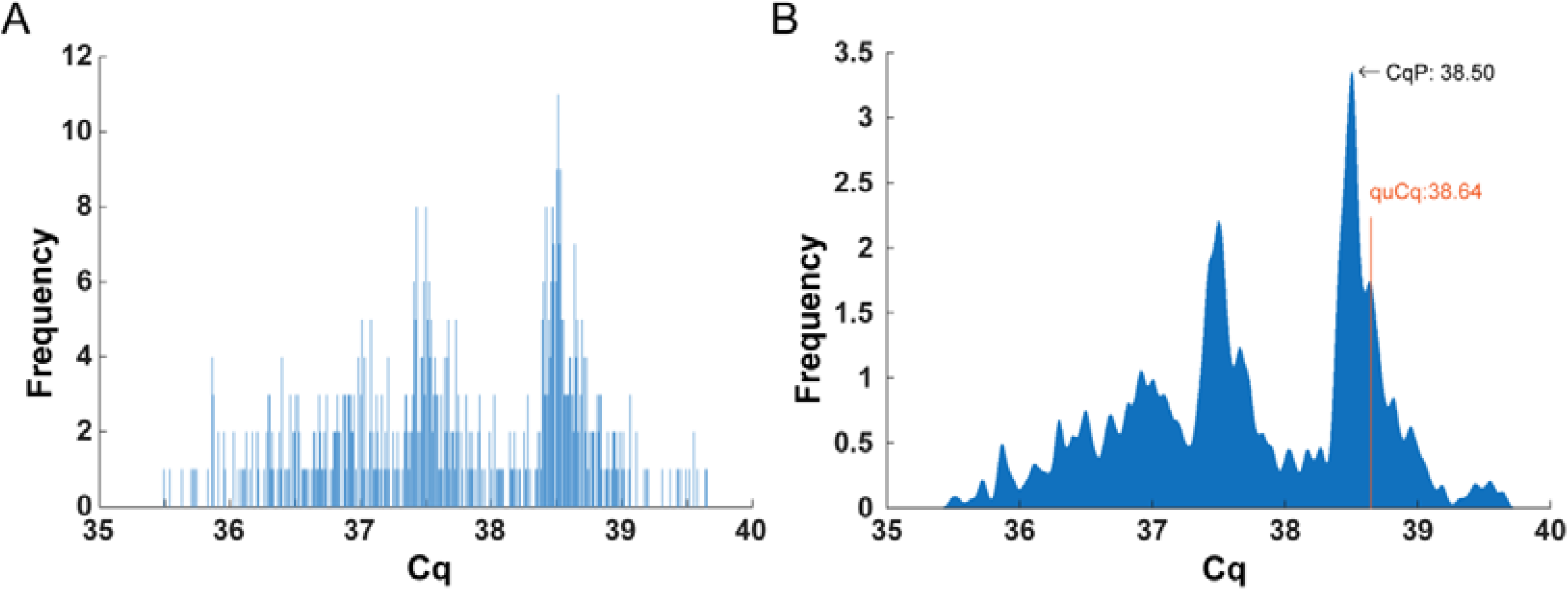
Cq distribution profiles for HER2-containing single copy ssDNA. **A)** The Cq distribution profile of 960 qPCR assays with a ssDNA template following limiting dilution of the template. The red curve shows the Gaussian-smoothed result.

**Table S1.**
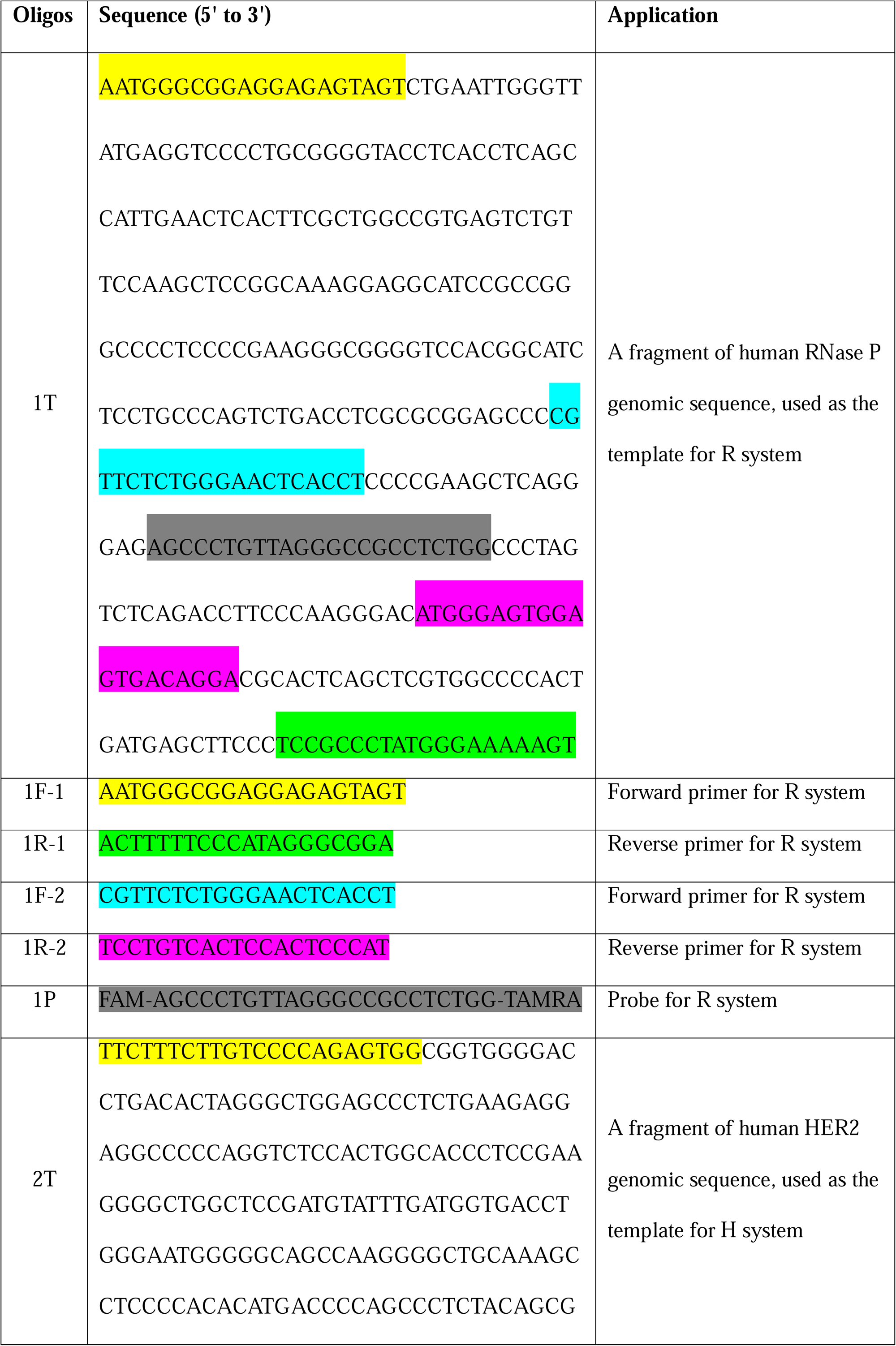
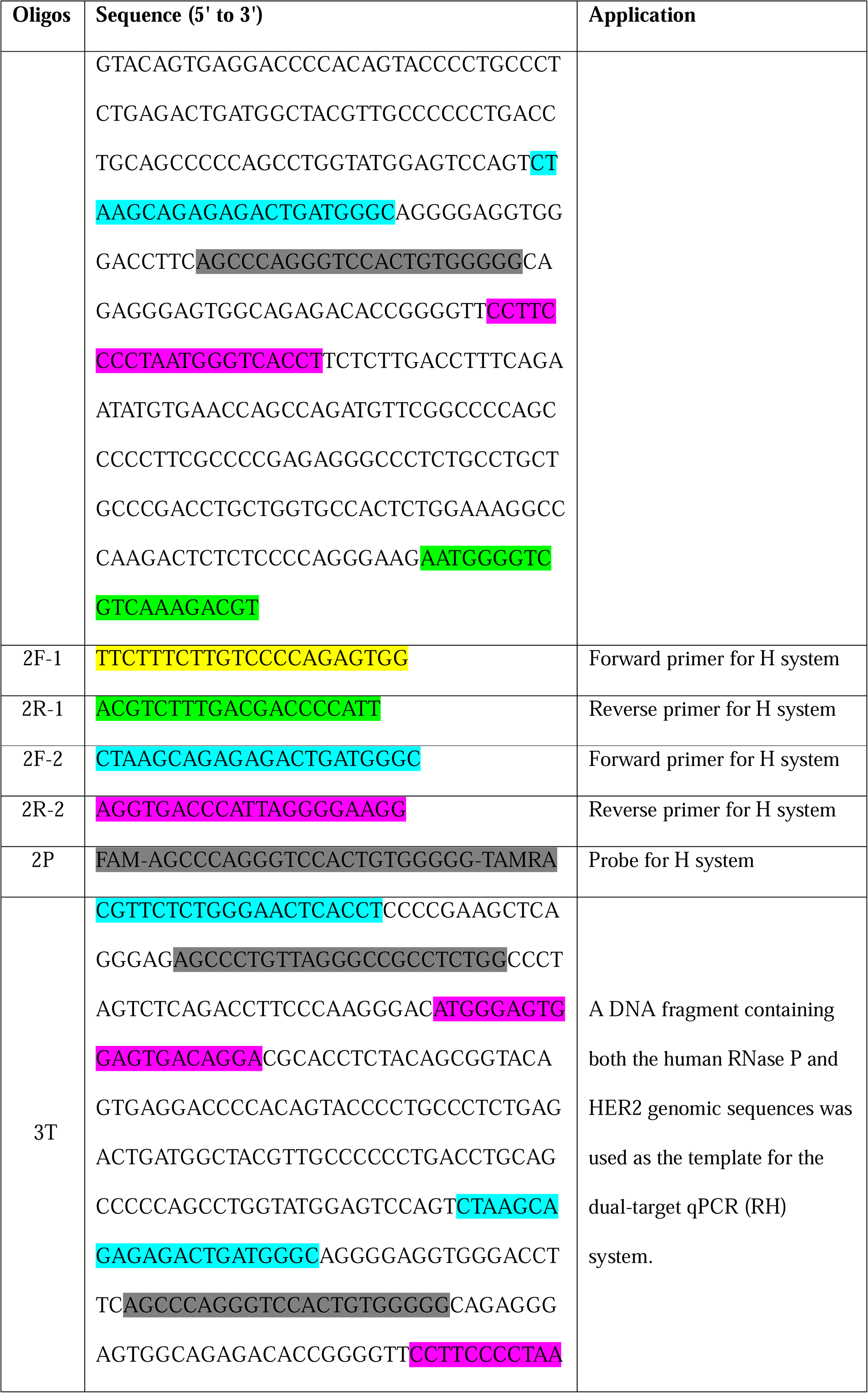
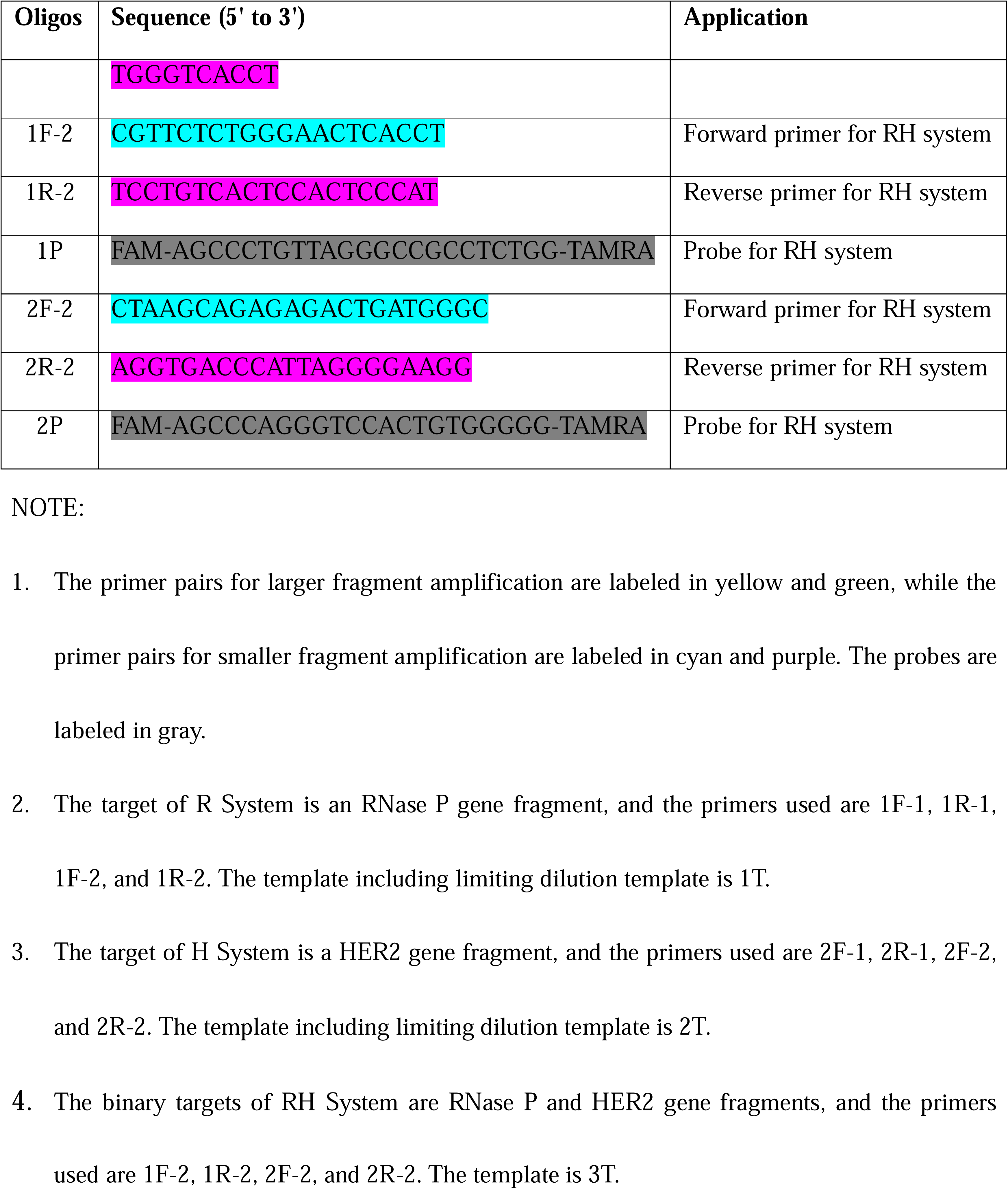
Oligonucleotides used in this study.

**Table S2.** Binomial distribution of sampling and dilution across multiple stocks.

| Initial copy | Dilution fold | Expected copy | Within 99% confidence interval |  |
| --- | --- | --- | --- | --- |
|  |  |  | Copy | Replicates needed for Average/Expected: (0.995-1.005) |
| 500000 | 50 | 10000 | 9745-10256 | 7 |
| 1000000 | 10 | 100000 | 99228-100773 | 1 |
| 500000 | 10 | 50000 | 49453-50547 | 2 |
| 50000 | 10 | 5000 | 4827-5173 | 14 |
| 5000 | 10 | 500 | 446-555 | 120 |
| 500 | 10 | 50 | 33-68 | 1327 |
| 50 | 10 | 5 | 0-11 | 11945 |

The binomial sampling error inherent in serial dilution. It quantifies the expected copy number variation and the minimum technical replicates required to ensure (with >99% confidence) that the mean measured concentration deviates by less than 0.5% from the expected value after dilution. The required number of replicates is calculated, for instance, diluting from 500,000 to 10,000 copies requires 7 technical replicates at least.

**Table S3.** Poisson distribution with various average copy numbers.

| **Average copy#**<br>**per well**<br>**Expected**<br>**copy# per well** | 0.5 | 1 | 1.5 | 2 | 2.5 | 3 | 4 |
| --- | --- | --- | --- | --- | --- | --- | --- |
| 0 | 60.65% | 36.79% | 22.31% | 13.53% | 8.21% | 4.98% | 1.83% |
| 1 | 30.33% | 36.79% | 33.47% | 27.07% | 20.52% | 14.94% | 7.33% |
| 2 | 7.58% | 18.39% | 25.10% | 27.07% | 25.65% | 22.40% | 14.85% |
| 3 | 1.26% | 6.13% | 12.55% | 18.04% | 21.38% | 22.40% | 19.54% |
| 4 | 0.16% | 1.53% | 4.71% | 9.02% | 13.36% | 16.80% | 19.54% |
| 5 | 0.02% | 0.31% | 1.41% | 3.61% | 6.88% | 10.08% | 15.63% |
| 6 | 0.00% | 0.05% | 0.35% | 1.20% | 2.78% | 5.04% | 10.42% |

The theoretical probability of finding a given number of template copies in a reaction well at different average concentrations ( $\lambda$ ) from 0.5 to 4. It was used to guide the dilution factor for the limiting dilution assay. Notably, across a broad range of average copy numbers from $\lambda=0.5$ to $\lambda=1.5$ — where the expected percentage of negative wells varies from about 60% to 20% — the theoretical proportion of wells containing exactly one molecule remains consistently around 1/3.

**Table S4.**
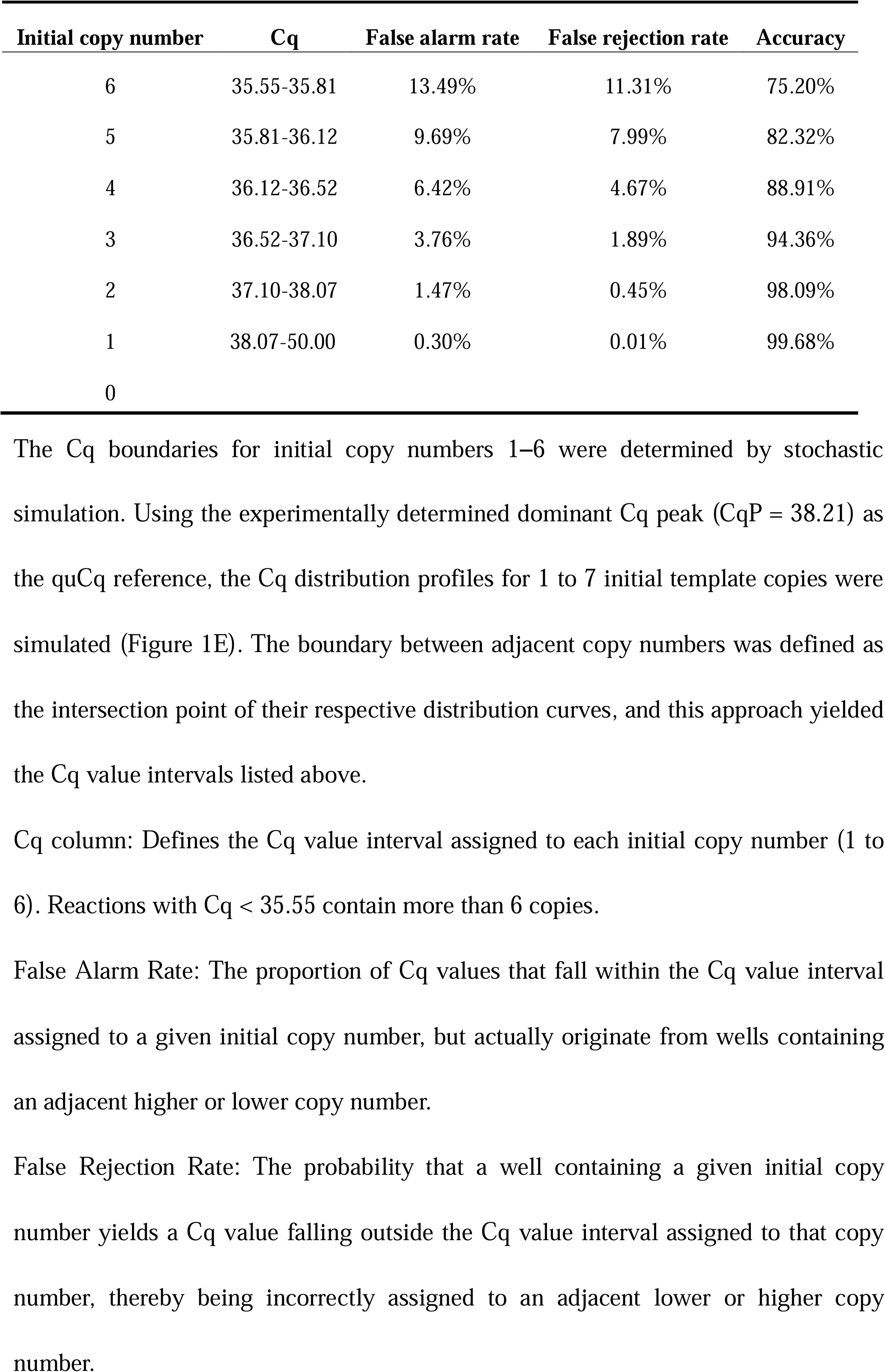
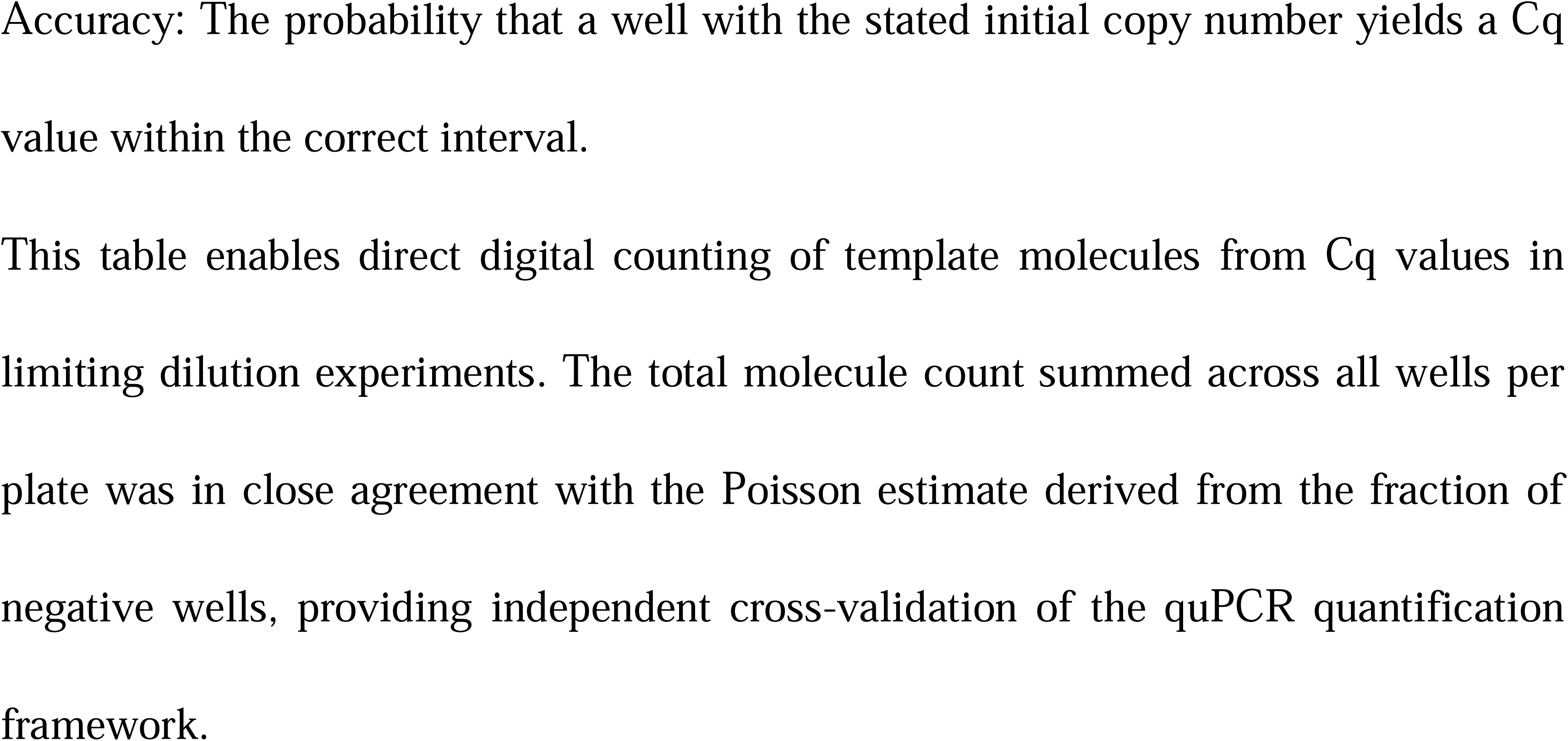
Correlation of Cq values with initial copy numbers.

**Table S5.** quPCR quantitation of binary targets and their ratio determination.

| Template | H Cq | R Cq | H copy | R copy | H/R | H/R | H/R |
| --- | --- | --- | --- | --- | --- | --- | --- |
| Dilution | (Average) | (Average) | numbers | numbers |  | (Mean) | (SD) |
| **1** | 19.08 | 18.90 | 490040 | 478137 | 1.025 |  |  |
| **2** | 21.46 | 21.27 | 99223 | 96995 | 1.023 |  |  |
| **3** | 23.83 | 23.62 | 20276 | 19935 | 1.017 | 1.022 | 0.035 |
| **4** | 26.23 | 25.94 | 4073 | 4209 | 0.968 |  |  |
| **5** | 28.62 | 28.47 | 824 | 765 | 1.078 |  |  |

H: HER2 gene, R: RNase P gene

